# DualMyo: Multi-Channel Dual-Stream Transformer Architecture for EMG-to-Digit Classification

**DOI:** 10.64898/2026.08.20.745897

**Authors:** Maria Golitsyna, Anna Makarova, Mikhail Lebedev

**Affiliations:** IAI MSU

## Abstract

Surface electromyography (sEMG) is a robust non-invasive modality for human-machine interaction, yet its application remains largely limited to coarse motor tasks such as grasping or rotation. The decoding of fine motor skills, specifically hand-writing, remains a challenging problem with potential relevance for prosthetic control and natural communication interfaces. In this work, we explore a Transformer-based alternative to classical signal-processing pipelines that treats multichannel sEMG signals as complex time series. We introduce DualMyo, a specialized model integrating Patch Embeddings and Rotary Positional Embeddings (RoPE) to capture the intricate spatiotemporal dynamics of myoelectric activity. Our experimental results show strong intra-session performance. Furthermore, we address the inherent challenges of signal drift and sensor displacement in cross-session applications. We show that a lightweight fine-tuning strategy of 10 epochs enables DualMyo to effectively adapt to session variability, achieving approximately 91% accuracy with two examples per digit. These findings provide a promising step toward adaptive sEMG-based hand-writing interfaces, although further validation is required for real-time and multi-subject deployment and neuromuscular control.

## 1 Introduction

Surface electromyography (sEMG) is a non-invasive technique for recording the electrical activity of muscles, which has found widespread application in biomechanics, rehabilitation medicine, and the development of human-machine interfaces. It is particularly valuable for prosthetics: the electrical signal from motor neurons is recorded on the skin surface milliseconds before the actual muscle contraction occurs. This predictive nature of the myoelectric signal allows the interface to detect the user’s motor intent prior to physical movement, potentially reducing control latency and enabling the natural control of bionic devices [5] [15].

Despite its high potential, the analysis of sEMG signals involves significant methodological challenges. The signal is highly susceptible to various artifacts and hardware noise. Recording quality critically depends on skin conditions (e.g., perspiration levels, body hair) and the reliability of electrode contact. During active movements, sensors can partially detach, generating anomalous noise spikes. Furthermore, electromyography is inherently non-stationary. Even when sensors are reapplied to the same anatomical zones of the same subject, the signal undergoes substantial changes. External factors such as ambient temperature, muscle fatigue, and even the subject’s psycho-emotional state (stress levels) significantly influence the characteristics of the recorded noise. This combination of factors leads to a severe domain shift between recording sessions. Consequently, models successfully trained on data from a single session often exhibit a sharp degradation in accuracy when transferred to new data without additional adaptation [15].

Historically, the majority of research in EMG interfaces has focused on decoding coarse motor patterns—such as clenching a fist, wrist rotation, or basic grasping gestures [5] [15] [2] [6] [22]. For such movements, the impact of noise is relatively minor, and numerous representative datasets are publicly available. However, the decoding of fine motor skills, specifically handwriting (e.g., classifying individual digits based on forearm muscle activity), remains a largely unexplored area. The myoelectric signals accompanying finger micro-movements possess low amplitude and high complexity, which may render classical approaches less effective for such low-amplitude and highly variable signals.

To address these challenges, we propose **DualMyo**, a compact Transformer-based architecture specifically designed for the classification of handwritten digits using sEMG signals. A defining feature of our model is its ultra-lightweight footprint—comprising only about 480,000 parameters—which makes it highly suitable for “on-device” training and inference on low-power edge devices.

Our approach differs from many existing approaches, which often rely on hard-coded, non-trainable signal filters or employ generic architectures that ignore the physiological specificities of EMG. A central design choice in DualMyo is its Dual-Stream architecture:

- **Raw Stream:** Extracts high-frequency components directly from the raw waveform using a trainable convolutional layer (PatchEmbed).
- **Residual Envelope Stream:** Computes the low-frequency signal envelope. Unlike the classical Root Mean Square (RMS) calculation, we apply the concept of Residual Learning. The model learns to adaptively correct a standard fixed envelope (env_fixed_) using a weighted combination of spectral basis bands (env_basis_).

This architecture, augmented with attention mechanisms, allows the model to capture both local high-frequency bursts and the global pattern of muscle activity without the quadratic increase in computational complexity typically associated with long sequence lengths.

### Contributions

In summary, the main contributions of this work are as follows:

1. We introduce **DualMyo**, a compact dual-stream Transformer for single-subject sEMG digit classification that combines channel-independent tokenization, RoPE-based temporal encoding, and gated fusion of raw-waveform and residual-envelope streams.
2. We evaluate DualMyo under intra-session and Leave-One-Session-Out cross-session protocols on six recording sessions of handwritten digit sEMG data, using trial-level splits to avoid window-level leakage.
3. We analyze the role of temporal context, channel-independent tokenization, and residual envelope modulation through cross-session experiments, few-shot adaptation, channel-contribution maps, and architectural ablations.
4. We show that DualMyo achieves high accuracy relative to the evaluated baselines in this single-subject setting, including 99.17% segment-level intra-session accuracy, 89.44% LOSO cross-session accuracy with 5,000-tick windows, and 91.22% accuracy after 10-epoch few-shot calibration with two examples per digit.

## 2 Related Work

The recognition of fine motor patterns through muscle activity is a critical task, not only for human-machine interfaces (HMI) but also for diagnosing neurological conditions affecting motor function. Consequently, researchers have long explored various methodologies to decode these signals.

In early studies, such as [14], classical algorithms like Linear Discriminant Analysis (LDA) and Principal Component Analysis (PCA) were proposed for digit classification, achieving intra-session accuracies of approximately 90%. Recent studies have focused on both traditional surface-based handwriting and the emerging task of “air-writing” [19]. In both domains, classical algorithms like Support Vector Machines (SVM) and k-Nearest Neighbors (KNN) have been extensively optimized, pushing intra-session accuracy to 95% [18], [17]. However, these methods rely heavily on manual feature engineering and remain highly sensitive to cross-session signal drift.

To address these limitations, deep learning approaches have been increasingly adopted. For instance, [3] evaluated hybrid architectures such as CNN-LSTM and CNN-GRU, which also achieved mean accuracies near 95% for handwriting tasks. Similar progress has been made in optical myography (OMG) [11], where models like TranScribe and GRUScribe were developed to predict writing trajectories. In the context of kinematic trajectory prediction, HandFormer [12] introduced a Transformer-based model capable of predicting 32 sequential hand positions, with each point represented by a 20-degree-of-freedom vector.

Furthermore, the proliferation of open-source datasets has led to the development of foundation models for physiological signal decoding [1, 4, 7–10, 13, 16, 20, 21]. Notable EMG-specific solutions include TinyMyo [8], EmgNet [13], and Physiowave [4]. While these models aim for universal EMG representations, they were not specifically trained on fine motor tasks, and their fundamental generalizability does not necessarily extend to handwriting decoding. Moreover, these architectures are significantly larger than our proposal, with parameter counts ranging from 1.75M (EmgNet) to 5M (Physiowave) and 3.6M (TinyMyo).

In contrast, our proposed **DualMyo** architecture achieves competitive accuracy in our experimental setting while maintaining a substantially smaller parameter count of only ∼480k parameters. To the best of our knowledge, few prior models have evaluated this task under a comparable parameter budget.

## 3 Methods

### 3.1 DualMyo Architecture Overview

DualMyo is a dual-stream Transformer architecture designed for cross-session electromyography (EMG) digit classification. Given an input EMG window **X** ∈ ℝ^*C×T*^ with *C* = 8 channels and *T* = 1000 samples, the model processes the signal through two complementary streams: a raw waveform stream and a residual envelope stream. The raw stream preserves the high-frequency morphology of the EMG waveform, while the envelope stream provides a smoother amplitude representation that is robust to local phase variations and session-dependent signal shifts.

### 3.2 Channel-Independent Tokenization

Both streams utilize the same channel-independent patching strategy. Each EMG channel is divided into non-overlapping temporal patches of length *L* = 10, resulting in *N*_*p*_ = *T/L* = 100 patches per channel and a total of *N* = *C* · *N*_*p*_ = 800 tokens per input window. Patches are embedded independently per channel using a 2D convolution with a kernel size of (1, *L*) and a stride of (1, *L*):

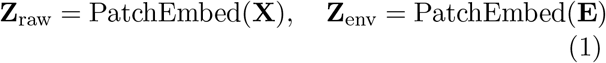

where **Z**_raw_, **Z**_env_ ∈ ℝ^*N ×*48^. The same learnable channel embedding is added to both streams, allowing the model to retain electrode identity while keeping temporal tokenization channel-independent.

### 3.3 Residual Envelope Stream

The envelope stream is built upon an adaptive residual envelope extractor. First, a fixed Root Mean Square (RMS) baseline envelope is computed per channel using a 50-sample moving window:

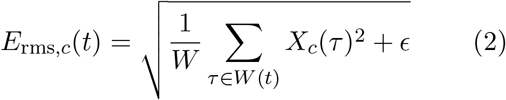

To allow the model to adapt this hand-crafted baseline, DualMyo concurrently computes a spectral envelope basis. The input signal is transformed via a Fast Fourier Transform (FFT), filtered by *K* = 8 fixed Gaussian band-pass masks with logarithmically spaced center frequencies within the 10–250 Hz range, transformed back to the time domain, and converted to RMS-style sub-band envelopes. These sub-band envelopes are mixed using learnable per-channel softmax weights:

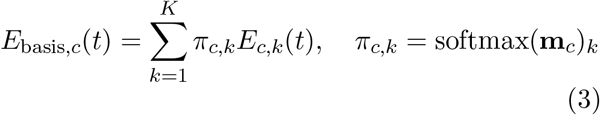

The final envelope is obtained as a bounded residual correction of the initial RMS baseline:

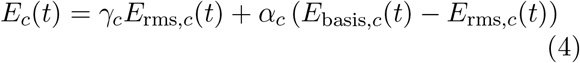

Here, the coefficient *α*_*c*_ = 0.2 tanh(*a*_*c*_) is initialized at zero and bounded to a small per-channel correction, ensuring the model initially stabilizes on the fixed RMS envelope. The scaling term *γ*_*c*_ = 1 + 0.2 tanh(*g*_*c*_) is initialized at one and bounded to the range [0.8, 1.2], enabling mild per-channel amplitude calibration. The output envelope is clamped to non-negative values.

### 3.4 Gated Dual-Stream Fusion

Following the patching layer, the raw and envelope token sequences possess matched lengths and dimensionalities. DualMyo fuses them token-wise using an adaptive gated fusion module. For each token, the raw embedding **r** and envelope embedding **e** are concatenated and passed through a sigmoid gate:

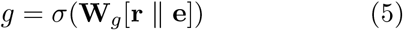

The fused token combines both information streams as follows:

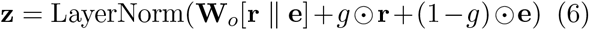

This formulation allows the network to adaptively emphasize either highly detailed waveform morphologies or macro-envelope dynamics depending on the specific channel, time segment, and class-discriminative feature pattern. The fused 48-dimensional tokens are subsequently projected to the Transformer embedding dimension *d* = 96.

### 3.5 Rotary Transformer Encoder

After gated fusion, the token sequence is projected from the per-stream fusion dimension to the Transformer hidden dimension *d* = 96. The resulting sequence contains *N* = *C* · *N*_*p*_ tokens, where *C* = 8 is the number of EMG channels and *N*_*p*_ = 100 is the number of temporal patches per channel. Importantly, tokens are ordered by concatenating all temporal patches of the first channel, followed by all patches of the second channel, and so on:

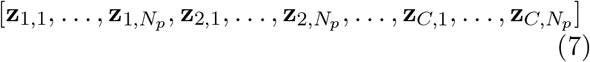

Therefore, tokens corresponding to the same temporal patch but different channels are separated by exactly *N*_*p*_ = 100 positions in the flattened sequence. For example, temporal patch *p* appears at flattened indices:

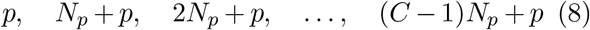

Because of this structure, we do not use the flattened token index as the positional coordinate. Instead, we assign RoPE positions according to the within-channel temporal patch index:

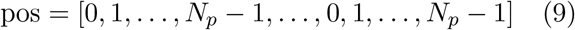

Thus, every *N*_*p*_-th token receives the same temporal position. This design encodes the assumption that tokens from different channels can correspond to the same time interval, while the learnable channel embedding identifies which electrode produced each token. In other words, temporal identity is encoded by RoPE, and spatial/electrode identity is encoded by the channel embedding.

Each Transformer block follows a prenormalization residual architecture:

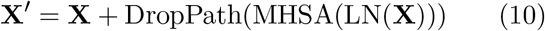

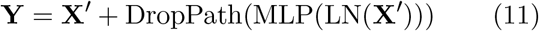

RoPE is applied inside the multi-head self-attention module to the query and key vectors only. For each attention head, the query and key representations are rotated according to their temporal patch position before computing scaled dot-product attention. This makes attention sensitive to relative temporal relationships between EMG patches without adding a learnable absolute positional embedding to the token vectors.

This is particularly suitable for the DualMyo token layout: tokens from different channels but the same temporal patch share the same RoPE phase, making cross-channel attention at aligned time points easy to learn, while attention between different temporal patches still receives position-dependent structure. As a result, the model can jointly reason over two axes: temporal dynamics through RoPE and inter-channel muscle activation patterns through self-attention and channel embeddings.

### 3.6 Classification Head

The encoder output is normalized and reshaped back into a channel-time structure. For each temporal patch, channel embeddings are concatenated:

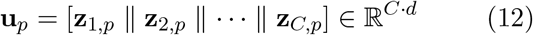

Temporal average pooling is then applied over the *N*_*p*_ patch positions, producing a single window-level representation. A linear classifier maps this representation to 10 digit logits. The classification head is intentionally lightweight and contains no hidden layers, ensuring that the bulk of the modeling capacity remains concentrated within the dual-stream tokenization, fusion, and the Transformer encoder layers.

## 4 Experimental Setup

### 4.1 Dataset and Evaluation Framework

The dataset used in this work comprises six independent recording sessions (*M* 1–*M* 6) of surface electromyography (sEMG) signals collected from a single subject at different points in time [14]. Within each session, myoelectric activity was recorded during the handwriting of 10 digits (0–9), with 10 repetitions performed for each character.

Biosignal acquisition was executed using an 8-channel sEMG sensor configuration. The electrodes were distributed anatomically to optimize the capture of diverse motor patterns: 4 sensors were positioned on the forearm, and 4 sensors were placed on the hand. This topology enables the simultaneous capture of macro-muscular contractions from the forearm alongside fine, localized phase variations of the hand muscles responsible for finger micro-movements during writing tasks.

To reduce temporal leakage, the intra-session 80/20% stratified split was performed at the trial level rather than at the window level. All windows extracted from the same gesture repetition were assigned exclusively to either the training or test partition, ensuring that neighboring windows from a single gesture did not appear in both splits.

### 4.2 Session-Level Signal Profiling

To characterize cross-session variability, we computed a small set of session-level signal statistics, including global RMS amplitude, trial-to-trial RMS variability, zero-padding rate, 50Hz power, channel-energy imbalance, and mean trial duration. These statistics were used only as descriptive diagnostics of session quality and recording variability; they were not used as model inputs. The resulting profiles are summarized in Table 1.

**Table 1:** Session-Level Signal Profiles Across Recording Sessions.

| Session | Global RMS | RMS std | Pad % | 50Hz Power | Channel Profile (Max/Min) | Mean Trial Length |
| --- | --- | --- | --- | --- | --- | --- |
| M1 | 0.9819 | 0.1343 | 1.88% | 0.0553 | Ch2: 20% / Min: 4% | 2240 ± 557 ms |
| M2 | 0.2925 | 0.0510 | 2.04% | 0.0048 | Ch0: 16% / Min: 7% | 2560 ± 767 ms |
| M3 | 0.3374 | 0.0670 | 2.95% | 0.0070 | Ch3: 17% / Min: 8% | 2304 ± 699 ms |
| M4 | 0.3815 | 0.0678 | 3.21% | 0.0085 | Ch0: 19% / Min: 7% | 2333 ± 654 ms |
| M5 | 0.3380 | 0.0619 | 0.48% | 0.0055 | Ch4: 21% / Min: 6% | 2364 ± 838 ms |
| M6 | 0.2291 | 0.0598 | 1.06% | 0.0023 | Ch5: 23% / Min: 6% | 2620 ± 667 ms |

The profiles indicate substantial variation across sessions. Sessions *M* 2–*M* 4 show relatively similar amplitude ranges, balanced channel-energy distributions, and low 50Hz power, whereas *M* 1 and *M* 6 show stronger deviations in amplitude and channel balance. Session *M* 5 occupies an intermediate regime, with low zero-padding but a shifted channel-energy profile. These differences motivate the use of cross-session evaluation and help contextualize the LOSO results reported below.

### 4.3 Baseline Configurations and Metrics

To benchmark the performance of the proposed **DualMyo** framework, we implement three baseline configurations spanning classical machine learning with handcrafted features and state-of-the-art deep bio-foundation models:

1. **Handcrafted TD + LogReg:** A traditional statistical learning baseline. We extract 6 standard time-domain (TD) features per channel: Mean Absolute Value (MAV), Root Mean Square (RMS), Standard Deviation (STD), Waveform Length (WL), Zero Crossings (ZC), and Slope Sign Changes (SSC). This yields a 48-dimensional feature vector from each *z*-score normalized input window. The classification is optimized using a grid-searched linear Logistic Regression classifier with L2 regularization.
2. **TinyMyo (Linear Probing):** Evaluated as a frozen foundation feature extractor. We utilize the state-of-the-art pretrained TinyMyo EMG backbone (8 layers, 192 hidden dimensions, ∼3.6M parameters) from the BioFoundation framework [8]. The pretrained weights are kept strictly frozen, and only a newly initialized 10-class linear classification head is optimized on the target session data.
3. **TinyMyo (Full Fine-Tuning):** Evaluated in an end-to-end optimization regime. Both the pretrained TinyMyo feature extractor backbone and the linear classification head are jointly updated, allowing all internal weights to adapt unconstrained to the target data distribution.

All baselines were evaluated using the same trial-level splits as the corresponding DualMyo experiments. TinyMyo and the handcrafted-feature baselines were trained and evaluated using 1,000-tick input windows.

### Inference Modalities and Evaluation Metrics

For our proposed DualMyo framework, we report performance across two primary inference modalities to evaluate temporal context processing:

1. **Window-Level Mode:** Inference is executed independently and non-contextually on isolated, discrete 1, 000-tick (or scaled 5, 000-tick) sample windows.
2. **Segment-Level Mode:** Predictions are temporally aggregated across a complete continuous gesture segment (the entire duration of writing a single digit) via mean probability voting across constituent windows prior to the final argmax operation.

Since the dataset is perfectly balanced across all ten digit classes (0–9) with exactly 10 physical repetitions per character within each recording session, standard top-1 **Classification Accuracy (%)** is adopted as the primary quantitative performance metric across all intra-session, cross-session, and ablation experiments.

### 4.4 Training Details

All DualMyo models were trained for up to 500 epochs with early stopping using a patience of 100 epochs. Optimization was performed using AdamW (torch.optim.AdamW). We used a batch size of 32, dropout of 0.1, label smoothing of 0.1, and weight decay of 10^−2^. The learning-rate schedule used cosine annealing with a maximum learning rate of 3 *×* 10^−4^, a minimum learning rate of 10^−5^, and 10 warmup epochs. Input preprocessing used per-channel normalization. During training, we applied the standard hard-augmentation pipeline used in our cross-session training script. This pipeline included Gaussian noise with standard deviation 0.02, random temporal shifts within *±*50 samples, channel dropout with probability 0.2, time masking with probability 0.4 over a 20–100-sample segment on a randomly selected channel, and crop-and-resample augmentation. These settings were kept fixed across the reported DualMyo experiments unless explicitly stated otherwise.

## 5 Results and Discussion

### 5.1 Intra-Session Decoding and Kinematic Error Dynamics

Under the intra-session decoding paradigm, where model training and evaluation are executed within the same recording session, the proposed **DualMyo** architecture achieves the best performance among the evaluated baselines compared to both traditional statistical feature pipelines and scaled foundation networks. Table 2 compiles the averaged classification metrics evaluated across all six independent sessions (*M* 1–*M* 6).

**Table 2:** Averaged Intra-Session Digit Classification Accuracy Across All Sessions (*M* 1–*M* 6)

| Model Configuration | Test Accuracy (%) |
| --- | --- |
| LogReg (Handcrafted 48d) | $94.40 \pm 4.81$ |
| TinyMyo (Linear Probing) | $60.97 \pm 12.65$ |
| TinyMyo (Full Fine-Tuning) | $86.70 \pm 9.61$ |
| **DualMyo (Ours, Window-Level)** | $97.18 \pm 1.94$ |
| **DualMyo (Ours, Segment-Level)** | **$99.17 \pm 1.25$** |

The benchmark results indicate that, even at the isolated window level, DualMyo yields an accuracy of 97.18%, outperforming the fully fine-tuned TinyMyo foundation model by over 10 pp. Crucially, introducing temporal aggregation at the segment level via probability voting enables DualMyo to substantially reduce the effect of local window-level variability within isolated windows, reaching 100.0% accuracy in three of the six sessions.

### Kinematic and Physiological Error Analysis

A detailed inspection of the predicted class probability distribution matrix (see Fig. 2) reveals potential kinematic and physiological patterns of the learned feature space. Rather than being distributed randomly, misclassifications can be structured and clustered tightly around two primary phenomena:

1. **Morphological Similarity (The Early Grapheme Effect):** A persistent mutual confusion is observed between digits “8” and “3”. In the biomechanics of handwriting, the initial kinematic trajectories of stroke formation for these two characters are identical. Because an isolated data window (especially under shorter temporal horizons) does not always encompass the full execution cycle of the symbol, the model relies heavily on this shared initial motor pattern. This observation suggests that future work could explore sub-symbolic or grapheme-level representations in future frameworks capable of dynamically reconstructing arbitrary textual sequences.
2. **Physiological Drift (Chronological Temporal Shift):** The error density distribution exhibits a distinct non-random tendency where, under high uncertainty, the network biases its predictions toward classes that are directly adjacent in the chronological recording sequence (e.g., misclassifying a “3” as a “4”). Given the sequential block-order of data collection during the experiments, this phenomenon suggests that the network may be sensitive to slow-varying physiological or recording-related factors.

**Figure 1:**
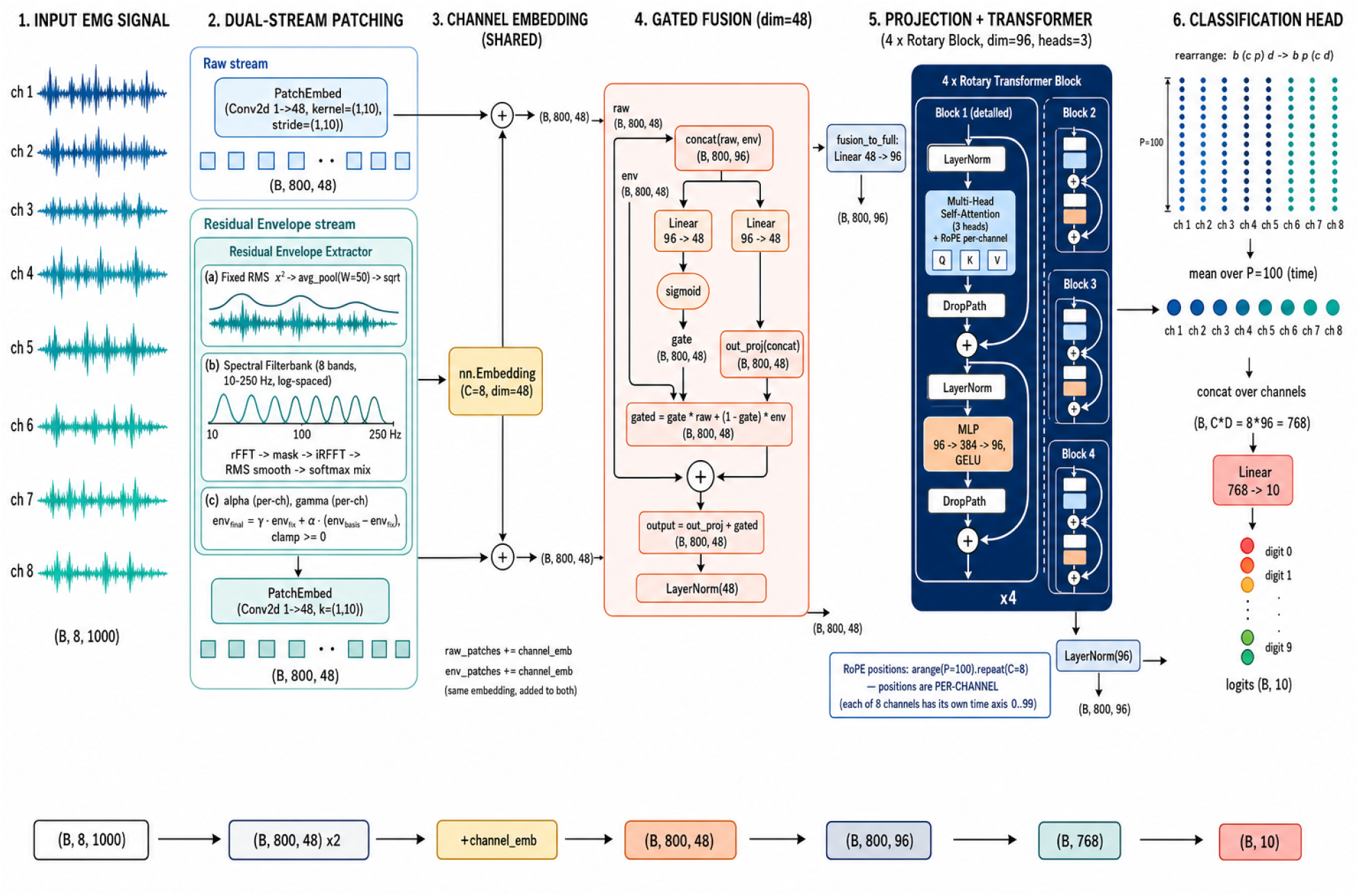
The DualMyo architecture overview, showcasing the Dual-Stream processing with the Raw Stream and the Residual Envelope Stream.

**Figure 2:**
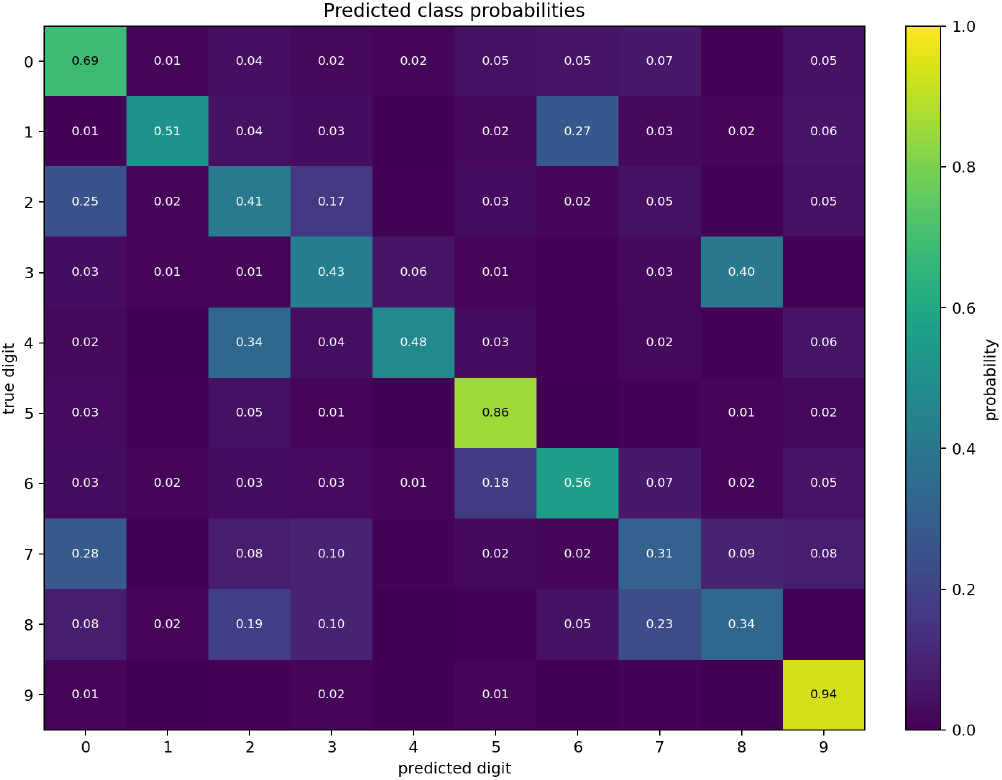
Class prediction probabilities for the intra-session model.

### 5.2 Cross-Session Generalization and Contextual Window Scaling

A stricter validation paradigm for modern sEMG interfaces is evaluating their robustness against cross-session domain shift. As shown by the statistical characterization in Section 4.2, phenomena such as physiological drift, alterations in skin-electrode impedance, and physical sensor displacement upon sensor re-application introduce severe non-stationarity into the underlying data distribution.

To quantify this factor under strict operational constraints, we implemented a Leave-One-Session-Out (*LOSO*) cross-validation protocol. Models were iteratively trained on a pool of 5 sessions and evaluated on the completely held-out 6th session. Table 3 compiles the final comparative metrics, with baseline accuracies aggregated across all independent cross-session test folds.

**Table 3:**
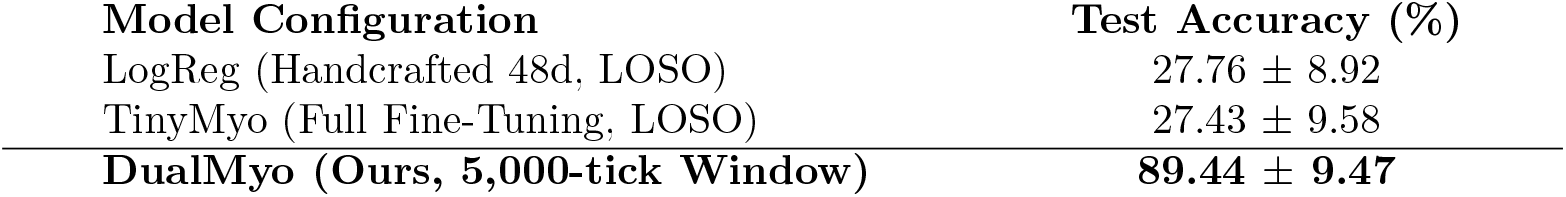
Averaged Cross-Session Digit Classification Accuracy Under Leave-One-Session-Out (LOSO) Validation.

#### Limitations of Foundation-Model Transfer in This Setting vs. Structural Inductive Biases

An inspection of the cross-session benchmarks in Table 3 reveals critical insights regarding model generalization:

1. **Limited Cross-Session Transfer of the Evaluated Foundation Model:** Despite being a scaled deep architecture pretrained on massive biomedical datasets, the *TinyMyo* foundation framework shows limited cross-session generalization in our evaluation, with an average accuracy of 27.43%. The fact that this deep foundation model matches the performance of a simple linear Logistic Regression over rudimentary time-domain features (27.76%) indicates that unconstrained end-to-end pretraining does not appear sufficient, by itself, to handle the sensor shifts present in this dataset. The network appears to overfit to the spatial coordinate systems of the training sessions and remains sensitive to shifted recording domains.
2. **Effect of DualMyo’s Channel-Independent Design:** Under identical conditions, our proposed DualMyo architecture achieves an average decoding accuracy of **89.44%**. By isolating channels at the tokenization level and deploying learnable sub-band envelope scaling, the network may encourage representations that are less sensitive to physical sensor displacement and layout misalignment.

#### Engineering Insight: The Role of Contextual Window Scaling

Empirical validation revealed that optimizing the temporal input horizon is statistically crucial for mitigating out-of-distribution drift.

When restricted to a baseline window of **1**,**000 ticks** (matching the intra-session setup), DualMyo yields an average window-level accuracy of 64.98%, though segment-level temporal voting recovers this to 79.86% (with localized improvements reaching up to +17.52% in stable domains like *M* 2). In this narrow temporal regime, localized windows contain insufficient macro-context, which may encourage the self-attention blocks to over-index on high-frequency session-specific noise.

Expanding the contextual tracking window to **5**,**000 ticks** substantially improved cross-session performance:

1. The model receives an extended temporal canvas that fully spans the execution envelope of the gesture, mapping the window-to-segment relationship effectively 1 : 1 and bypassing the need for downstream voting heuristics.
2. This scaling was associated with improved performance in distorted recording blocks: the spatially shifted session *M* 5 increases from 68.67% to **89.07%**, while the severely attenuated session *M* 6 is improved from 20.04% to **80.32%**.

Consequently, macroscopic window scaling may act as an implicit regularizer by providing more complete gesture context, helping the model rely more on global movement structure than on local session-specific variations.

### 5.3 Physiological Interpretability and Gated Fusion Analysis

The token-wise gated fusion mechanism integrated into DualMyo provides a possible interpretation of channel contributions. The gating values and contribution maps offer a way to inspect how the model uses different channels. The feature decomposition curves for individual digits (Fig. 3) and the cumulative channel impact map (Fig. 4) illustrate a structured distribution that is broadly consistent with the electrode layout.

**Figure 3:**
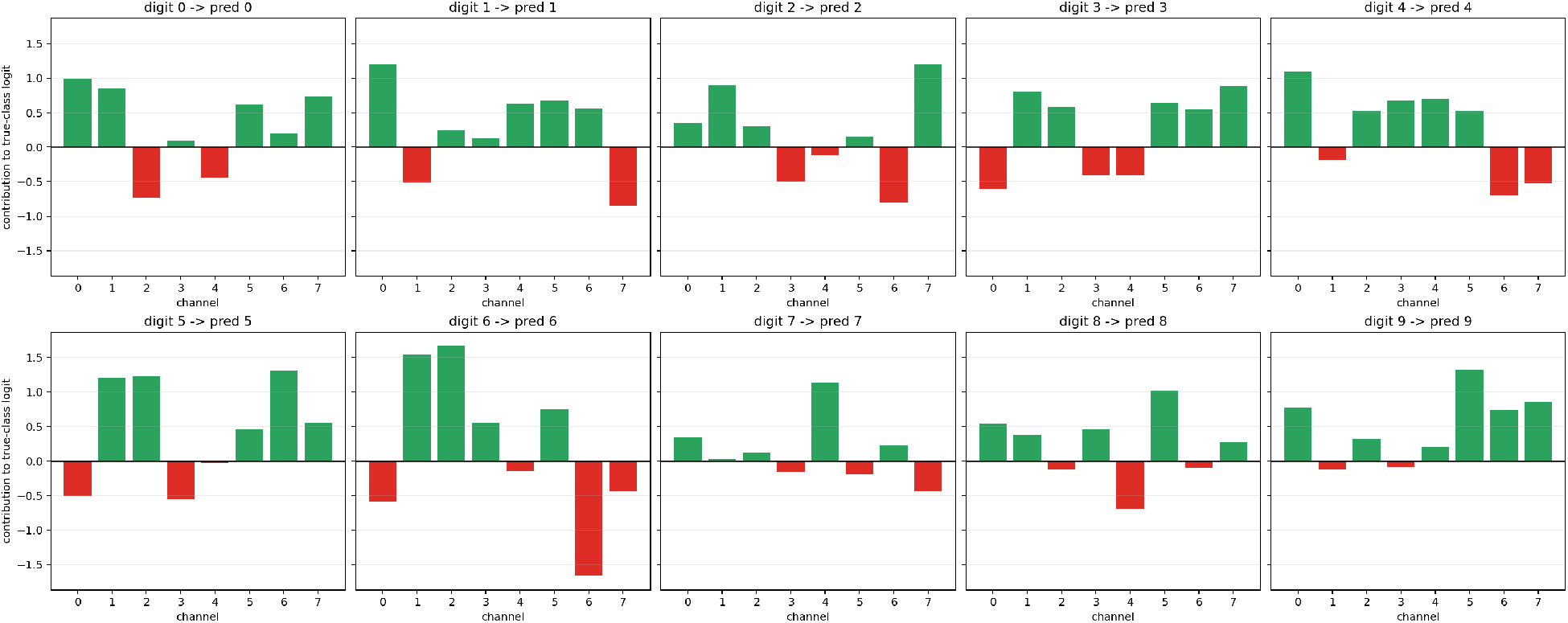
Contribution of EMG channels to class prediction.

**Figure 4:**
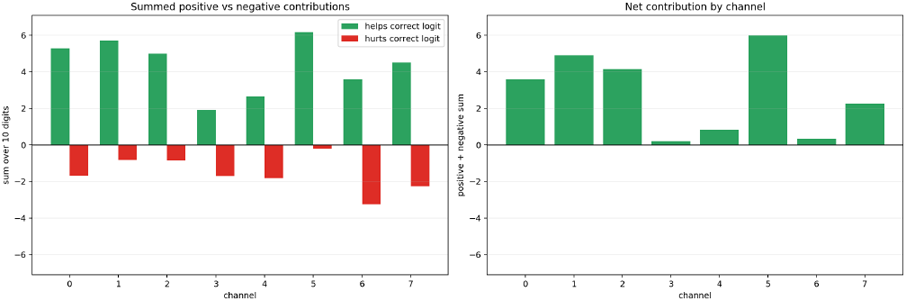
Aggregate contribution of EMG channels to class prediction.

#### Channels with Larger Positive Contributions (Forearm Group)

The largest positive model contributions are observed for sensors located on the forearm (Channels 0, 1, and 5). Anatomically, Channels 0 and 5 map onto the superficial projections of the *m. extensor carpi radialis* and *m. extensor carpi ulnaris* (the radial and ulnar wrist extensors), while Channel 1 corresponds tightly to the *m. flexor carpi radialis* (the radial wrist flexor). The observed contribution pattern is consistent with the hypothesis that wrist and hand stabilization contribute important information for digit classification.

#### Fine Motor Modulation (Hand Group)

Micro-movements and stroke adjustments are captured by the sensors distributed across the hand (Channels 2, 3, and 4). Specifically, Channel 3 is positioned near the thenar region, which may explain its contribution. This channel shows a relatively consistent positive contribution across nearly all ten digit classes, reflecting its sensitivity to the fine-grained thumb micro-adjustments executed during complex pen stroke transitions.

#### Selective Channel Suppression

Channels 4 and 6 show a distinctive behavior in the interpretability profiles. Channel 4, which topologically shares the thumb thenar region alongside Channel 3 (mapping near the *m. abductor pollicis brevis*), yields a near-zero net aggregate contribution. Concurrently, Channel 6 shows negative logit contributions across several digit classification tasks.

This behavior suggests that DualMyo may learn to suppress less informative or noisy channels. Instead, through its learnable gated fusion routing (Equations 5 and 6), the network can be interpreted as learning a form of adaptive spatial weighting. It may down-weight less informative channels while assigning larger positive contributions to channels associated with more informative motor patterns.

#### Benefits of Channel-Independent Tokenization

This operational capacity highlights why channel-independent tokenization performed better than early spatial channel mixing in our ablation study (such as the 1*×*1 convolution explored in the experiment with early channel mixing). Early channel mixing combines raw voltage signals before independent sub-band envelopes are computed. This may reduce the spatial specificity of individual muscles and spread power-line noise or electrode artifacts across clean channels. Strict channel isolation through the tokenization stage preserves channel-specific information that can be used by self-attention for non-linear spatial weighting.

### 5.4 Few-Shot Adaptation and Calibration Efficiency

In practical human-computer interface (HCI) deployments, requiring an end-user to undergo extensive data collection to initialize or recalibrate an electrode layout is highly undesirable. A viable system should support rapid adaptation, maximizing performance gains from a small calibration set. To quantify the few-shot adaptation capacity of the **DualMyo** architecture, we systematically evaluated its cross-session fine-tuning dynamics under constrained data regimes and limited optimization horizons.

Our experimental protocol simulates a small-calibration paradigm on a completely held-out test session under two few-shot data limits: *n* = 1 (where the user records each digit from 0 to 9 exactly once) and *n* = 2 (two physical repetitions per character). To reflect deployment-oriented constraints on low-power edge hardware, the fine-tuning window was strictly bounded to a 10-epoch horizon (requiring less than a few seconds of compute). These bounded runs are contrasted against both the uncalibrated zero-shot baseline (*Pre-FT*) and the best observed accuracy under unconstrained fine-tuning (*Max Acc*).

We benchmark these metrics across two distinct pre-training validation pipelines:

1. **With Validation (Strict 6-Session Core):** A rigorous validation pipeline where one session within the pre-training pool is fully isolated as a validation set for early stopping. This ensures zero target domain look-ahead and a completely objective checkpoint selection prior to fine-tuning on the hidden target domain.
2. **Without Validation (5-Session Pool):** A pipeline where the model is pre-trained across a combined pool of 5 sessions without an early-stopping validation set, and fine-tuning check-points are selected directly against peak performance on the target session.

Table 4 summarizes the comparative performance across these evaluation paradigms.

**Table 4:** Fine-Tuning Efficiency and Calibration Robustness Across Few-Shot Regimes.

| Evaluation Metric | With Validation (%) | Without Validation (%) | $\Delta$ (pp) |
| --- | --- | --- | --- |
| Pre-FT Per-Trial Accuracy | 84.02 | 89.44 | +5.02 |
| Accuracy @ Epoch 10 ($n = 1$) | 79.06 | 86.00 | +6.94 |
| Accuracy @ Epoch 10 ($n = 2$) | 85.00 | 91.22 | +6.22 |
| Max Accuracy ($n = 1$) | 86.98 | 90.41 | +3.43 |
| Max Accuracy ($n = 2$) | 91.09 | 93.61 | +2.52 |

#### Analysis of Validation Bias and Selection Non-Stationarity

While the metrics for the configuration “Without Validation” are systematically higher across all evaluation points in Table 4, it is important to highlight that this comparison does not represent an “apples-to-apples” scenario due to two confounding factors:

1. **Optimistic Selection Bias:** The “Without Validation” pipeline selects its optimal pretraining checkpoints by evaluating directly on the target test session. This introduces an implicit domain look-ahead bias that can lead to optimistic accuracy estimates.
2. **Anomalous Session Exclusion:** Crucially, the 5-session pool utilized in the unvalidated mode completely excludes session *M* 6 (export1 24Jan08). As established in the statistical data profiling (Section 4.2), *M* 6 represents a strong out-of-distribution session characterized by substantial signal attenuation and possible electrode-contact degradation. Conversely, the strict “With Validation” protocol naturally absorbs the severe non-stationarity of *M* 6 within its cross-validation folds.

Despite encountering significantly harsher data distributions and zero domain look-ahead, the strictly validated model shows substantial robustness. In the *n* = 2 maximum accuracy regime, the performance gap between the strict validation setup and the idealized unvalidated setup narrows to just 2.52% (91.09% vs. 93.61%).

More importantly, under the highly restricted 10-epoch limit with only two calibration samples (*n* = 2), the strictly validated model recovers immediately to a per-trial accuracy of **85.00%**. This rapid convergence suggests that the architecture can adapt efficiently under limited calibration. DualMyo’s channel-independent tokenization and residual envelope routing may act as a useful regularizing bottleneck, allowing the Transformer to adapt swiftly to complex physical sensor displacement without overfitting to high-frequency localized noise.

### 5.5 Architectural Ablation Study

To rigorously isolate the driving factors behind DualMyo’s generalization capacity and validate our core inductive biases, we conducted a broader ablation study and report representative configurations in Table 5. All ablation variants were bench-marked under the cross-session Leave-One-Session-Out (*LOSO*) paradigm on the designated target domain (export1 15Jan08), mapping performance deltas (Δ) in percentage points (pp) against the optimized baseline configuration. Note that the accuracy of our baseline model reported here is lower than the values presented in the previous sections, as it is evaluated strictly on a single target session using isolated 1,000-tick window-level samples without any temporal aggregation across trials.

**Table 5:** Architectural Ablation and Preprocessing Exploration Results.

| Explored Configuration / Hypothesis | Acc (%) | $\Delta$ (pp) |
| --- | --- | --- |
| **DualMyo Baseline (Learnable $\gamma + \alpha$)** | **57.79** | — |
| *Phase 1: Multi-Stream and Modulation Variants* |  |  |
| Triple-Stream (Raw + Envelope + Highpass) | 48.48 | -9.31 |
| Triple-Stream (Raw + Envelope + Delta) | 53.45 | -4.34 |
| Gamma-only Modulation (No Filterbank Residual) | 49.45 | -8.34 |
| Envelope Delta FiLM Modulation | 53.66 | -4.13 |
| *Phase 2: Spatial Filtering and Signal Hygiene* |  |  |
| Common Average Reference (CAR) Filter | 49.31 | -8.48 |
| Quality Control: Drop Artifact-heavy Trials | 52.90 | -4.89 |
| *Phase 3: Sensor Augmentation and Capacity* |  |  |
| Augmentation: Unconstrained Circular Channel Shift | 46.76 | -11.03 |
| Capacity: Frozen $\gamma$ Parameter Calibration | 49.17 | -8.62 |
| Capacity: Reduced Hidden Layer Scale ($D = 96, L = 2$) | 46.07 | -11.72 |
| *Phase 4: Tokenization Boundary Analysis* |  |  |
| Early Spatial Channel Mixing (Pre-Patch $1 \times 1$ Conv) | 33.45 | -24.34 |

#### Importance of Channel-Independent Tokenization

The most severe performance degradation occurred when introducing early spatial mixing prior to temporal patching (a 1 *×* 1 Conv1d layer), which precipitated a large performance drop of −24.34 pp. This empirical result indicates that this component is important for performance in our setting. Prematurely blending multi-channel raw voltage waveforms may obscure fine-grained spatial differences and localized phase variations that the subsequent self-attention blocks utilize to separate macro-muscular contractions from finger micro-movements.

#### Anatomical Spatial Priors vs. Unconstrained Augmentation

A common regularization practice in standard circumferential muscle-sensing rings is to apply unconstrained data augmentations, such as random circular channel shifting (*±*1), to simulate sensor rotation. However, applying this technique to our architecture induced a severe performance drop (−11.03 pp). Crucially, our sensor topology is not an equidistant, symmetric ring; it maps 4 sensors directly to the forearm and 4 sensors to the hand to capture structurally distinct motor patterns. Consequently, unconstrained circular shifting violates fixed anatomical priors. Spatial channel swapping is mathematically valid only when bounded within localized, functionally symmetric physiological groups that align with the specific electrode placement.

#### Effects of Spatial Filtering and Data Cleaning

Counter-intuitively, standard preprocessing choices did not improve performance:

1. **Common Average Reference (CAR):** Implementing CAR severely damaged decoding accuracy (−8.48 pp). Subtracting the instantaneous global mean across all electrodes effectively strips away key spatial contrast and micro-amplitude variations required to decipher fine handwriting kinetics, suggesting that the loss of localized spatial information may outweigh the benefit of common-mode noise rejection.
2. **Data Cleaning:** Explicitly removing or down-weighting artifact-heavy or noisy trials consistently reduced cross-session generalization. This may indicate that the model benefits from exposure to realistic physiological noise during training. Training the network on idealized, perfectly “cleaned” trials may hinder it from learning domain-invariant representation spaces required to navigate real-world inter-session drift.

#### Optimal Feature Capacity and Bounded Modulation

Attempts to expand the architecture into a triple-stream configuration by adding raw high-pass or delta derivatives increased model complexity without improving performance and introduced optimization instabilities. Conversely, restricting the network’s envelope calibration by freezing the *γ* scalar reduced the model’s ability to perform mild amplitude normalization, inducing a −8.62 pp penalty. This suggests that our current dual-stream design with a bounded learnable spectral residual (*α, γ*) provides a favorable trade-off in our experiments—compact enough to avoid overfitting, yet sufficiently flexible to absorb inter-session sensor displacement.

## 6 Limitations

This study should be interpreted within the scope of a single-subject, multi-session evaluation. Although the recording sessions exhibit substantial non-stationarity and sensor-placement variation, the present experiments do not establish how well the observed trends transfer across subjects with different anatomy, writing styles, skin-electrode properties, or muscle activation patterns. Multi-subject validation is therefore required before making broader claims about general-purpose sEMG handwriting decoding.

The dataset is also limited to digit classification with a small number of repetitions per class. This setting is useful for studying fine motor decoding and cross-session robustness, but it does not yet cover continuous text entry, letters, punctuation, or naturalistic writing conditions. As a result, the reported performance should be viewed as evidence for digit-level decoding under controlled conditions rather than as a complete end-to-end myoelectric text interface.

Finally, our evaluation is performed offline. The deployment-oriented claims are motivated by the compact model size and short fine-tuning horizon, but real-time experiments are still needed to measure latency, user adaptation, electrode reapplication effects, and calibration burden in an interactive setting. Future work should therefore extend the evaluation to multi-subject recordings, online inference, and longer gesture sequences.

## 7 Conclusion

In this work, we investigated the problem of fine motor skill recognition through the lens of surface electromyography (sEMG) decoding during continuous digit handwriting. Our analysis exposed the fundamental constraints of existing large-scale biomedical foundation models, such as TinyMyo, when adapted to high-precision kinematic tasks.

Despite being pre-trained on massive, generalized datasets, these architectures showed limited transfer under cross-session domain shifts, possibly because their pre-training data may not capture the fine motor patterns required for this task. This suggests the value of exploring specialized models that integrate tight structural constraints and physiological signal priors.

Our proposed dual-stream architecture, DualMyo, improves cross-session generalization in this dataset through structural and algorithmic integration of domain knowledge. Specifically, isolating the signal amplitude envelope into a dedicated parallel processing stream, coupled with learnable per-channel scaling parameters (*α, γ*), may encourage the network to construct robust latent representations. As a result, DualMyo shows strong stability: it achieves near-perfect performance in intra-session scenarios (99.17% on a per-trial basis) and preserves high accuracy relative to the evaluated baselines of 89.44% under rigorous cross-session testing before target-session calibration in the evaluated LOSO setting. Furthermore, if a rapid deployment optimization is introduced, the architecture exhibits promising few-shot adaptation performance, climbing to 91.22% within a constrained 10-epoch calibration horizon and reaching up to 93.61% when optimization limits are removed.

A key engineering insight from our empirical evaluation is the critical role of the temporal input horizon. The experiments indicate that sEMG classification models may benefit from operating over longer windows that capture more complete gesture trajectories, such as the full trajectory of a written symbol captured within a 5,000-tick window. Attempting to chop the continuous data stream into isolated, short intervals (e.g., 1,000 ticks) heavily distorts the contextual landscape and substantially reduces cross-session generalization in our experiments.

This finding outlines a highly promising direction for future research, transitioning from isolated classification toward continuous, end-to-end myoelectric text decoding. Since segmenting full words or sentences into independent alphabetical characters can increase classifier complexity and cumulative error rates, a more logical evolution lies in decomposing handwriting gestures into graphemes—the elementary geometric and biomechanical building blocks of writing. Developing a compact, universal dictionary of graphemes rather than expanding the alphanumeric character set could reduce data collection bottlenecks and may enable downstream recurrent or autoregressive decoders to map continuous sEMG signal trajectories directly into contiuous text streams in real time.

